# Tracing social mechanisms and interregional connections in Early Bronze Age Societies in Lower Austria

**DOI:** 10.1101/2025.02.10.636471

**Authors:** Anja Furtwängler, Katharina Rebay-Salisbury, Gunnar Neumann, Fabian Kanz, Harald Ringbauer, Raffaela Bianco, Tanja Schmidt, Lena Semerau, Rita Radzeviciute, Rodrigo Barquera, Nadin Rohland, Kristin Stewardson, J. Noah Workman, Elizabeth Curtis, Fatma Zalzala, Kim Callan, Lora Iliev, Lijun Qiu, Olivia Cheronet, Anna Wagner, Guillermo Bravo Morante, Michaela Spannagel, Maria Teschler-Nicola, Friederike Novotny, Domnika Verdianu, Ron Pinhasi, David Reich, Johannes Krause, Philipp W. Stockhammer, Alissa Mittnik

## Abstract

In this study, we present the results of archaeogenetic investigations of Early Bronze Age individuals from Lower Austria, specifically associated with the Únětice and Unterwölbling cultural groups. Through analysing newly generated genome-wide data of 138 individuals, we explore the social structure and genetic relationships within and between these communities. Our results reveal a predominantly patrilocal society with non-strict female exogamic practices. Additionally, Identity-by-Descent (IBD) analysis detects long-distance genetic connections, emphasizing the complex network of interactions in Central Europe during this period. Despite shared social dynamics, notable genetic distinctions emerge between the Únětice and Unterwölbling groups. These insights contribute to our understanding of Bronze Age population interconnections and call for a nuanced interpretation of social dynamics in this historical context.

## Introduction

The transition from the Final Neolithic to the Early Bronze Age in central Europe during the third millennium BCE was marked by significant socio-cultural transformations and population dynamics. The migration of people from the Pontic-Caspian Steppe into Central Europe on an immense scale reshaped the genetics of Central European populations (**Lazaridis et al. 2014, Haak et al. 2015, Allentoft et al. 2015**). This migration led to encounters of the new coming pastoralists with sedentary farmers, in parallel with the widespread adoption of metallurgy. The advent of bronze as a valuable commodity played an important role in trade networks and fostering the exchange of goods, knowledge, and cultural practices as well as genes among diverse communities (**Fokkens and Harding 2013, Stockhammer and Maran 2017**).

Studies across different regions of Central Europe have documented compelling evidence for gradual genetic homogenization among populations during the Early Bronze Age (EBA). These studies reveal striking patterns of genetic similarity and shared ancestry among EBA individuals across large geographic areas, underscoring the extensive genetic interconnectedness across large geographic areas (**Mittnik et al. 2019, Furtwängler et al. 2020, Papac et al. 2021, Penske et al. 2024**). Moreover, these EBA populations exhibit notable similarities in settlement patterns and subsistence strategies, characterized by homesteads comprising residential and farming structures close to cemeteries.

Previous research has shed light on the social structure prevalent during this period and the preceding Copper Age, indicating patrilocal communities where sons inherited the family home while females married into other groups (**Mittnik et al. 2019, Sjögren et al. 2020, Penske et al 2024**).

These insights of largely homogenous and patrilocal EBA populations in Central Europe were mainly derived from an interregional perspective. However, the question remains how these dynamics manifested within smaller, more localized key regions.

To address this question, we focused on Lower Austria north of the Eastern Alps, a region with rich archaeological evidence from the Early Bronze Age, covering a discrete area of 80 kilometers in radius. The area is divided by the river Danube, which gave rise to the formation of two distinct early Bronze Age cultural expressions in immediate vicinity: the Únětice and the Unterwölbling groups, which differ significantly in their burial custom.

The area north of the Danube is part of the Unětice complex, known for its archaeological treasures such as the famous Nebra Sky Disc and for social stratification expressed in early ‘princely’ burials such as those at Leubingen and Helmsdorf in Central Germany (**Meller 2017, Pernicka *et al*. 2020**). In contrast, the archaeological evidence from Lower Austria is more modest **(Lauermann 2003)**. It consists of small farmsteads, often near rivers and bodies of water, with settlement burials in former storage pits, and cemeteries of a few dozen graves near the settlements. The graves are sometimes arranged in rows in small plots, and while a single burial per grave is commonplace, instances of double and multiple burials are also present. The predominant burial practice involved positioning both male and female bodies in a flexed position, typically on their right side, oriented with the head in the south and the gaze directed towards the east. Archaeologically preserved grave goods are modest and comprise bone, shell and bronze jewelry and dress items, as well as ceramic bowls, jugs and vessels placed with the body. Tools and weapons such as daggers are rare. Animal bones and botanical remains from offerings of food and drink may also be present. Our archaeogenetic investigation focused on the archaeologically and anthropologically well-contextualized sites of Drasenhofen **(Horváth 2019, Neumann et al. 2023**), Zwingendorf (**Wewerka 1982**), Unterhautzenthal (**Rebay-Salisbury *et al*. 2018**), Schleinbach (**Pany-Kucera *et al*. 2020)** and Ulrichskirchen (**Verdianu 2024**) (see supplementary S1 for more information).

The Unterwölbling Group south of the Danube and west of the Viennese woods culturally connects to the Early Bronze Age southern German groups (**Neugebauer 1994**). Among the well-known sites from this region are the farmsteads and cemeteries from the Lech river valley, which have been extensively researched (**Knipper *et al*. 2017, Mittnik *et al*. 2019**). Following Bell Beaker traditions, and in contrast to the Lower Austrian Unětice groups, the burial rites were strongly gendered, even for children (**Neugebauer 1991, Rebay-Salisbury et al. 2022**). Male individuals were typically placed in a flexed position on their left side with their head to the north, while female individuals were usually positioned on their right side with their head to the south. Both thus faced towards the east. Gendered burial was practiced in large parts of Central Europe, for example in the southern Rhine area, along the Danube, as well as along large parts of the Tisza and Vistula rivers (see **Meller 2021**: 104 for a map). It appears that it was important to the Bronze Age people in these communities to differentiate between women and men. However, the way individuals were buried in gendered burials does not necessarily reflect how they identified themselves, highlighting the potential complexity of gender identity in these societies. Significant cemeteries have been discovered, with hundreds, if not thousands, of burials, such as at Gemeinlebarn F (**Neugebauer 1991**) and Franzhausen I and II (**Neugebauer and Neugebauer 1997, Reiter 2008**). Single burials are typical, and double and multiple burials are exceedingly rare. The depth of graves and the number and quality of grave goods vary according to the social status of the deceased, although this is difficult to determine due to the frequent reopening of graves when grave goods were removed. In this archaeogenetic study, we included individuals from Franzhausen I (**Neugebauer 1991**) and Pottenbrunn (**Blesl 2006**) (see supplementary S1 for more information).

By focusing on the microscale perspective, we aim to determine the level of genetic homogeneity across populations separated by geographically restrictive barriers, identify differing social structures within contemporaneous populations in Lower Austria, and enhance our understanding of the intricate genetic, social and cultural interactions within the EBA societies of Central Europe.

## Results

### General sample overview

We screened 198 prehistoric individuals from seven archaeological sites dating between 2300 BCE and 1600 BCE (see SI table 1, 2 and 7 and Fig. 1A and 1C) and applied a stringent selection process to ensure the quality of preserved ancient human DNA. Specifically, we enriched 143 individuals for 1,233,013 ancestry informative sites using the “1240k capture panel” (**Mathieson et al. 2015**) of the human genome.

**Figure 1:**
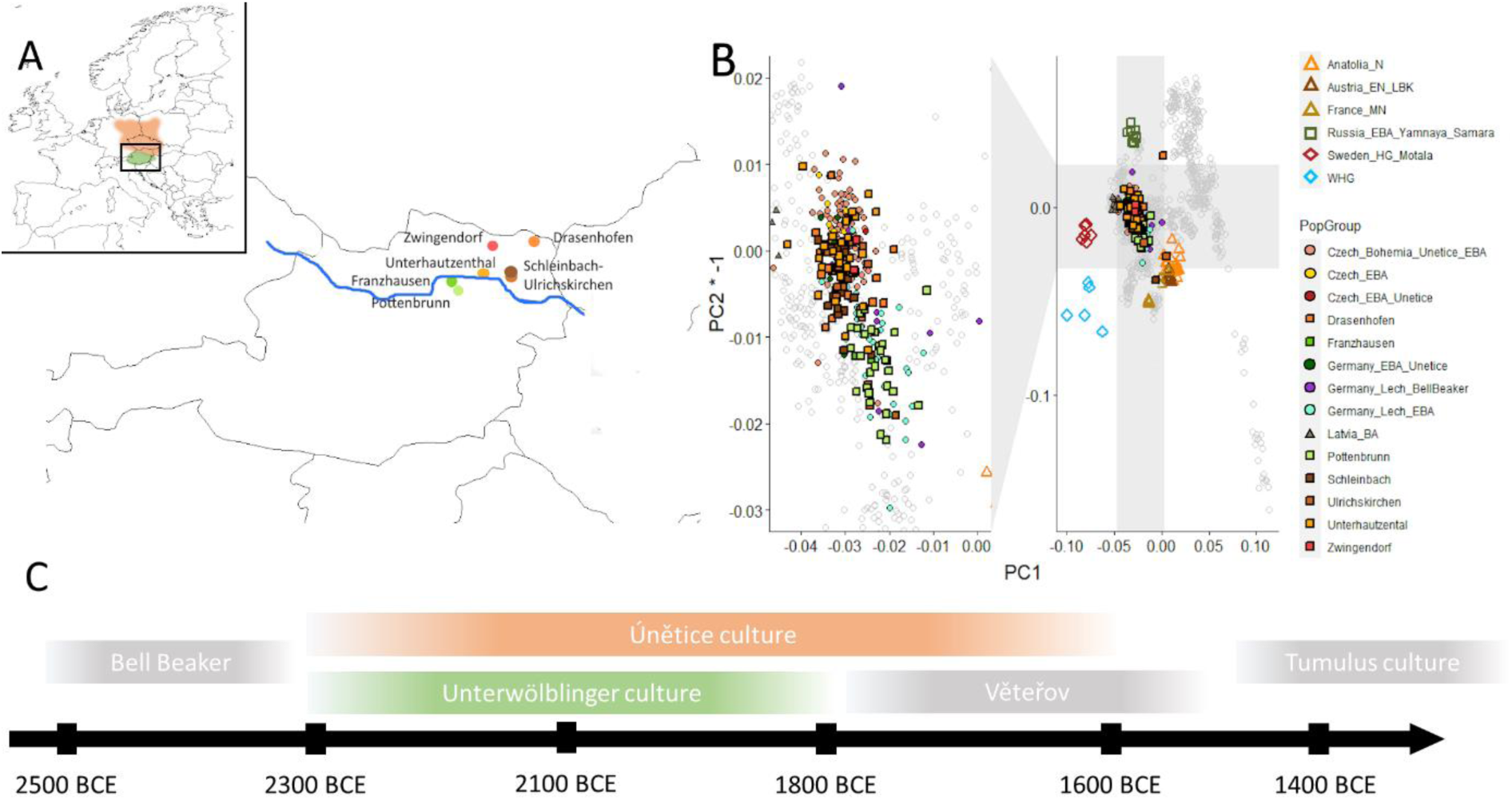
Temporal and geographic distribution of studied Early Bronze Age individuals from Lower Austria. A) Map of Austria showing the locations of sampled sites associated with the Únětice (orange) and Unterwölbling (green) cultures. B) PCA of modern West-Eurasian populations (grey) with projected ancient genomes. C) Temporal range of the Únětice and Unterwölbling culture in Lower Austria.

Following enrichment, individuals with insufficient coverage (less than 30,000 covered sites on the 1240k panel) or indications of contamination (more than 5% estimated contamination on the mtDNA and/or X chromosome) were excluded from the analysis, resulting in the removal of eight samples. As a result, we present a newly reported dataset comprising 138 individuals from the Early Bronze Age in Lower Austria.

### Genetic analysis of EBA groups in Lower Austria

We initially conducted a principal component analysis (PCA) by projecting the ancient individuals from EBA Lower Austria onto the first two axes. This PCA was constructed using a dataset of 551 modern-day West Eurasian individuals from 67 groups. In addition, we included 267 previously published ancient genomes (**Mathieson et al. 2015, Feldman et al 2019, Mathieson et al 2018, Olalde et al. 2018, Olalde et al. 2018, Furtwängler et al. 2020, Narasimhan et al. 2019, Jones et al. 2015**) as reference points and projected them onto the PCA plot alongside the EBA individuals from Lower Austria.

The resulting PCA plot (Fig. 1B) revealed distinctive patterns. As expected, all 138 EBA individuals from Lower Austria clustered between Early Farmers from Anatolia (Anatolia_Neolithic), Western Hunter-Gatherers (WHG), and pastoralists from the Pontic Caspian Steppe (Russia_EBA_Yamnaya_Samara) and exhibited close proximity to previously published contemporaneous groups from central Europe (**Olalde et al. 2018, Mittnik et al. 2018, Mittnik et al. 2019, Papac et al. 2021, Patterson et al. 2021**).

The genetic analysis indicates genetic differences between individuals from the areas north and south of the Danube. These differences are evident through the mean values observed on the second principal component (PC2) of the previously described PCA (Fig 1B and Fig 2A), as well as in the relative proportions of ancestry components modeled with *qpAdm* (Fig. 2B, SI table 3). Specifically, the individuals which inhabited the area south of the Danube (carrier of Unterwölbling culture) from Pottenbrunn and Franzhausen I exhibit a higher relative amount of Early Farmer ancestry in comparison to their Steppe-related ancestry. Conversely, the Únětice individuals individuals north of the Danube display the opposite pattern, with a greater proportion of Steppe-related ancestry relative to Early Farmer ancestry resulting in a ratio of Yamnaya Samara to Anatolia Neolithic of 1.91 for the Únětice and 0.89 for the Unterwölbling samples. These contrasting genetic profiles indicate significant genetic differences between the two populations, emphasizing the distinct ancestral contributions and genetic dynamics that influenced them. Analysis of runs of homozygosity (ROH, SI Note 3) indicates that both groups had a large and outbreeding population (as seen from SI Fig. 2) with no indication of unions between consanguineous parents.

**Figure 2:**
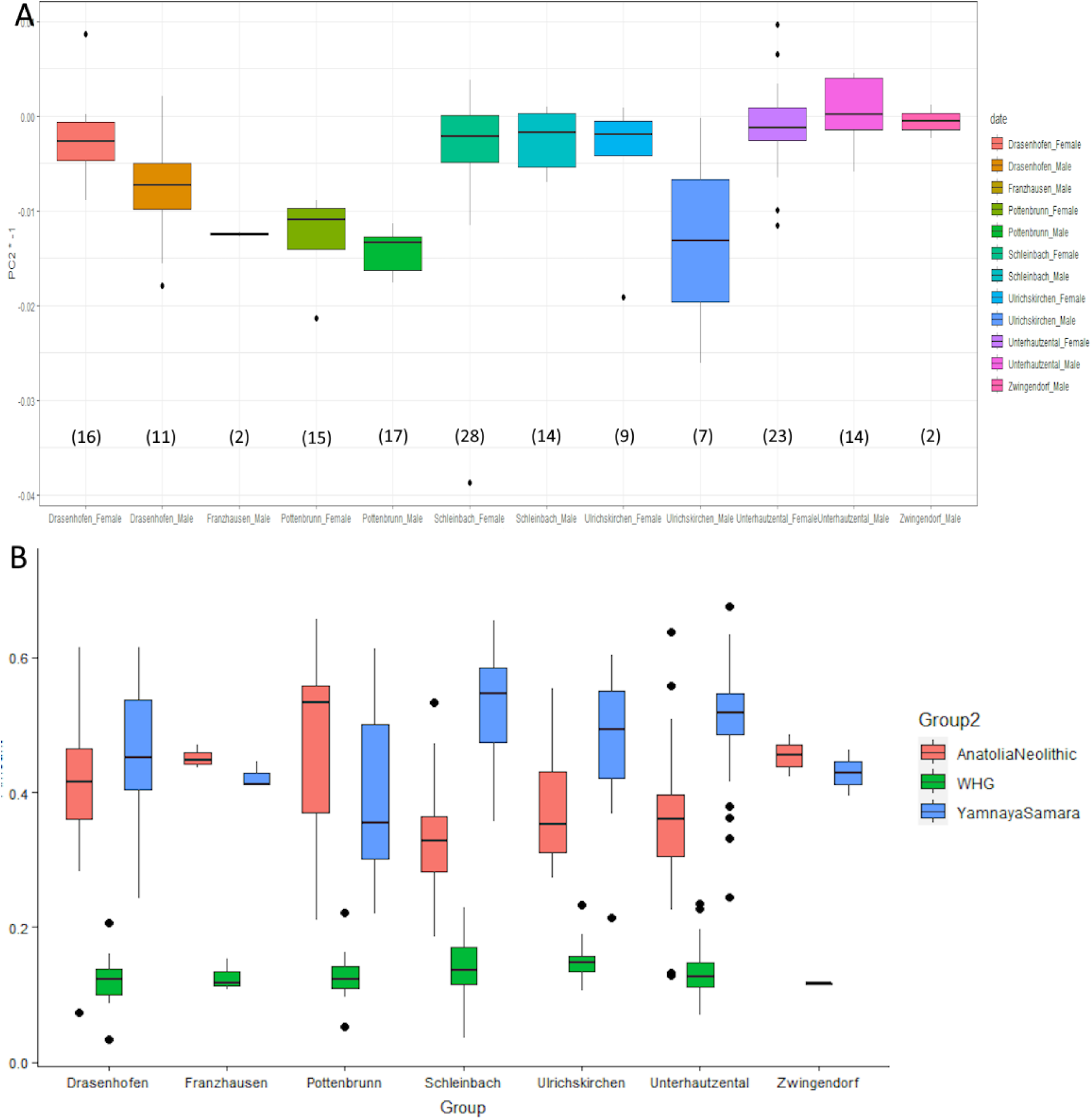
Differences in the mean values of PC2 and the relative amounts of estimated ancestry components for the Únětice and the Unterwölbling associated individuals. A) Mean values of PC2 separated by site and genetic sex, number in brackets indicate sample size. B) Relative amounts of ancestry components estimated with *qpAdm* for each individual grouped by site with mean values indicated by lines.

**Figure 3:**
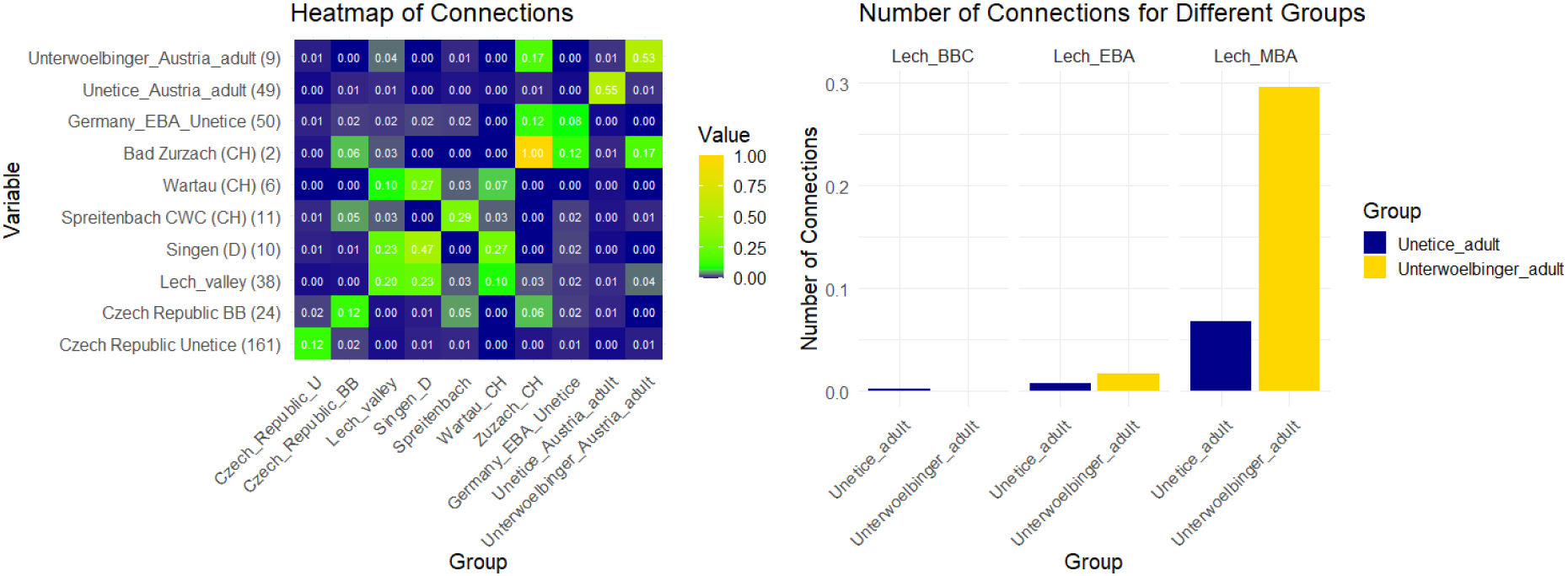
Comparison of IBD Connection Patterns Across Different Groups. A) Heatmap illustrating the number of IBD connections od segment length 8 cM, 12 cM, 16 cM and 20 cM normalized by the number of possible connections between various groups. The number of possible connections is calculated by n1×n2n between groups and n(n−1)/2 within groups following the method suggested by **Ringbaur et al 2024**. The values range from blue (low connection) to yellow (high connection), indicating the strength of the connections. B) Bar plot showing the normalized number of IBD connections for different groups in the Lech valley, categorized by Únětice and Unterwölbling. The x-axis represents different groups, while the y-axis shows the number of IBD connections. The colors distinguish between the Únětice and Unterwölbing categories.

When examining a set of functional SNPs (SI table 6), which includes lactose persistence in adulthood, there were no frequency differences observed between the two groups. The frequency of SNPs associated with lactose tolerance was generally low in total 18% of all investigated EBA individuals from Lower Austria carry one or two allels associated with lactase persistance.

### Intercultural connections between individuals associated with the Únětice and Unterwölbling culture

The investigation utilizing Identity-by-Descent (IBD) analysis allows the detection of genetic connections up to the 8th degree and even identifies connections that extend beyond that range (**Ringbaur et al. 2024**). The analysis of contemporary groups from different regions reveals that the studied individuals from the two cultures generally have the most genetic connections to the Lech Valley, specifically during the Middle Bronze Age (MBA). This pattern holds true even when considering only the Únětice culture. However, focusing solely on the Unterwölbling individuals, the strongest genetic connections are found with the Early Bronze Age (EBA) double burial from Bad Zurzach in Switzerland (**Furtwängler et al. 2022**).

IBD analysis furthermore reveals minimal connections between the two cultural groups under study even though all sites in this study originate from an area with no more than an 80-kilometer radius. Only one subadult individuals from the Unterwölbling sites Franzhausen display IBD segments shared with individuals from the Únětice group (Fig. 4, separate networks for males and females in SI note 5, SI table 5), albeit beyond the 8th degree of kinship. One adolescent boy from the triple burial of Franzhausen. He was buried with an adult man and a second adolescent boy who are identified as father and son, to which he is not genetically related.

**Figure 4:**
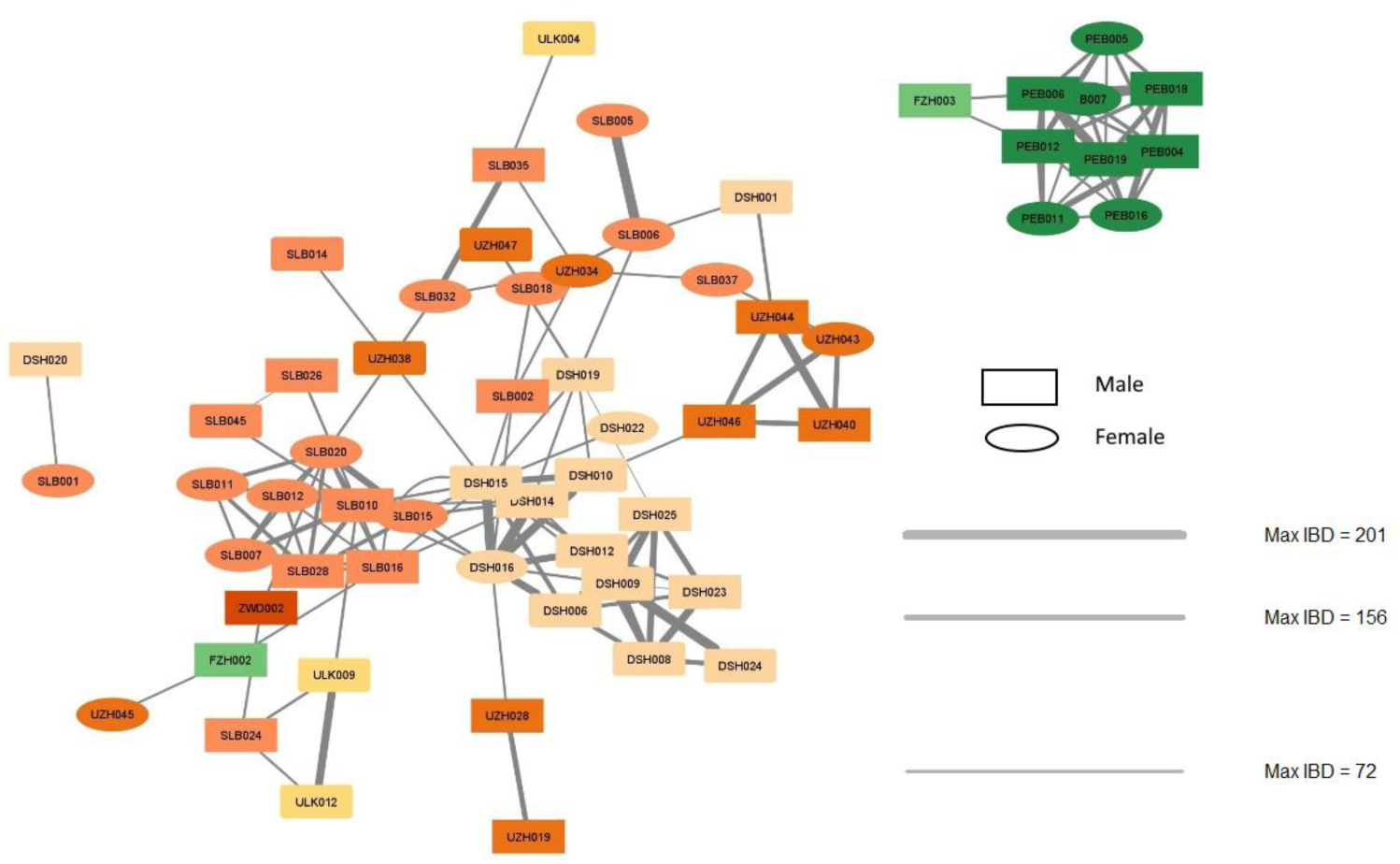
IBD sharing within and between the sites associated with the Únětice and the Unterwölbling culture. For individuals associated with the Unterwölbling culture in green and the Únětice culture in orange, the thickness of the lines indicated values of max IBD. Rectangles represent male individuals and ovals represent female individuals.

The IBD analysis also reveals notable gender-based differences in social connectivity within Bronze Age cultures, measured by the degree centrality (*k*), defined as the number of links held by each node. In the Únětice culture, females average degree centrality ⟨*k*⟩ is 2.54, compared to 3.42 in males. Dividing the number of links within site (*k_w_*) by the total number of links for each Únětice individual results in an *k_w_/k* ratio of 0.46 for females and 0.58 for males. This further shows that males are, on average, more connected within their site than females. Conversely, in the Unterwölbling culture, females average 5 links compared to 5.6 in males. Higher link values in males suggest a tendency towards patrilocality (**Gnecchi-Ruscone et al. 2024**).

### Genetic kinship relations within cultures and social structure

By integrating multiple methods, including the examination of mitochondrial DNA (mtDNA) and Y chromosomal haplogroups, we conducted a comprehensive analysis of genetic kinship and successfully reconstructed family trees within burial grounds associated with the two cultural groups (Fig. 5, SI table 4). These family trees extended up to three generations, revealing patterns of genetic relationships. It is important to note that genetic relatedness does not necessarily represent social kinship.

**Figure 5:**
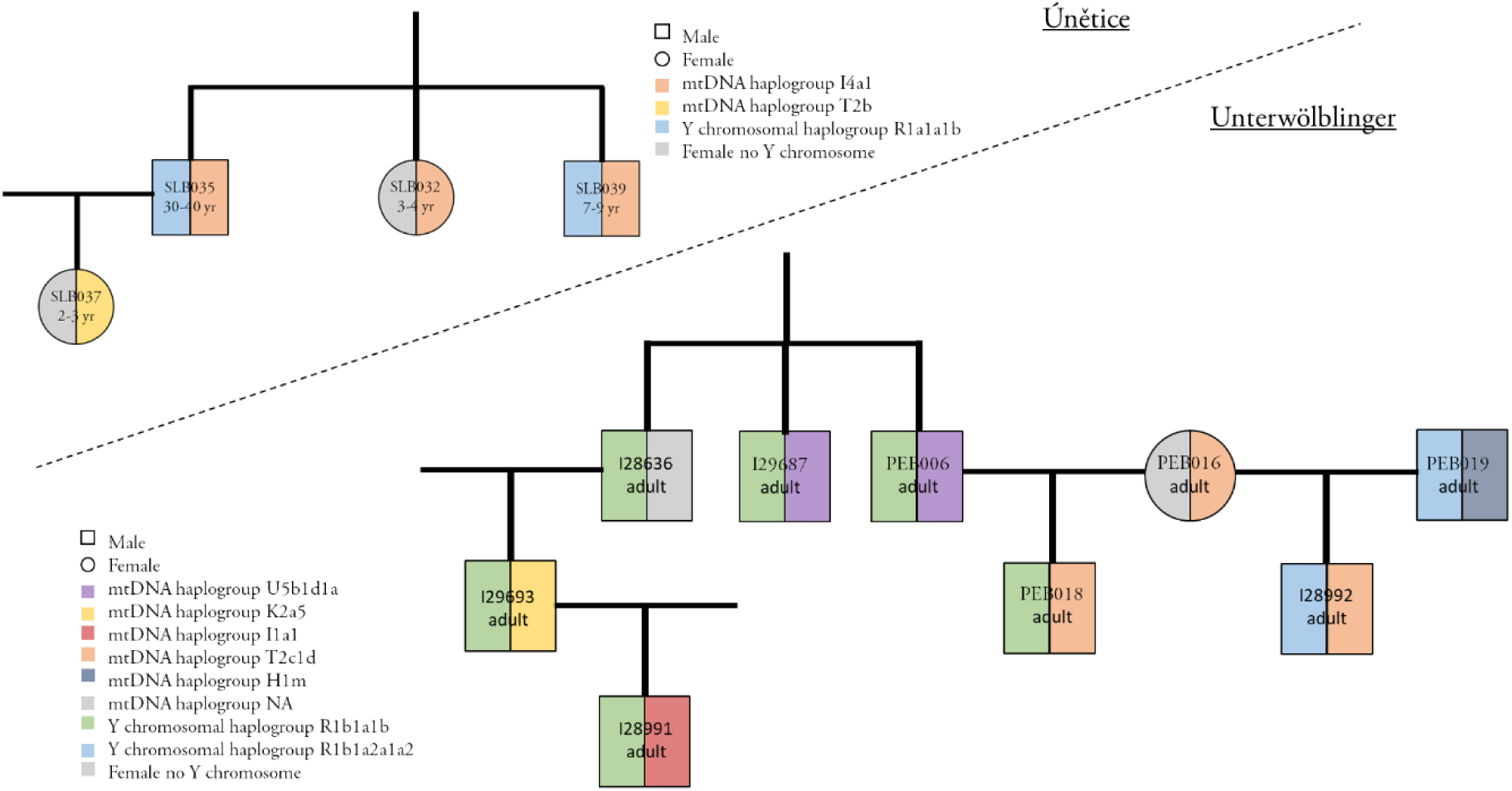
Examples of reconstructed Family Trees for the Únětice and Unterwölbling associated sites. This figure illustrates family trees from the Early Bronze Age Únětice culture (above the line) and the Unterwölbling culture (below the line). In each diagram, the left half of the circles and squares represent the Y chromosomal haplogroup, while the right half indicates the mtDNA haplogroup. Both cultures exhibit similar patrilocal residence patterns, with women primarily situated within their husband’s family units.

Nevertheless, our observations suggest that there was a common practice among males to be buried in the same cemetary as their parents and offspring, which might indicate a continuation of familial connections. In contrast, females were more commonly buried solely with their offspring.

In one family tree (SI Fig. 4) there are three siblings, two full-siblings with one half-brother who shares the same father, who was not among the sampled individuals. Because DSH023 and DSH027 share X chromosome similarities (SI table 4), this may suggests that the previous partner (DSH027’s mother) and DSH009, the mother of DSH008 and DSH027, share a distant common ancestor. However, the number of shared SNPs on the X chromosome is limited.

Our findings also unveiled two instances in the Únětice group where adult women were buried with their parents or siblings (SI Fig.3 and SI Fig. 10), which was unexpected based on previous research. In other regions of Central Europe during the Early Bronze Age, evidence of female exogamy was prevalent (**Mittnik et al. 2019**). One of these women, buried at Drasenhofen also has offspring buried in the same burial ground, indicating that also biological mothers were involved in these exceptions to female exogamy. These cases of adult women buried alongside close relatives suggest a more nuanced social dynamic, potentially indicating unique familial and social relationships within the studied populations North and South of the Danube.

## Discussion

The findings from this study present a notable advancement in our understanding of the Early Bronze Age (EBA) communities in the Danube region, as the near-exhaustive sampling of individuals for nuclear DNA from the burial grounds allowed for the construction of a rich and comprehensive dataset. This dense sampling has provided an unprecedented opportunity to gain a detailed snapshot of the people inhabiting this specific area during this prehistorical period.

Despite their geographical proximity, significant genetic differences exist between the Unterwölbling and Únětice groups. The individuals associated with the Unterwölbling culture exhibit a higher genetic ancestry related to the first farmers from Anatolia, in contrast to the higher proportion of steppe-related ancestry observed in the Únětice individuals. This pattern counters the geographical prediction and implies that the process of population homogenization in Central Europe during the EBA was driven by culturally shaped interconnectivity and trading networks, and was not a blanket phenomenon. Rather, these nuanced genetic distinctions point towards diverse regional dynamics, where different cultural groups may have chosen specific regions to foster close connections and facilitate the movement of individuals, particularly women. The significant genetic similarities on the X chromosome between half-brothers with the same father suggest that these females may come from the same region, as it implies that the foreign women were distantly related.

The strong connections of the Unterwölbling group to Bad Zurzach in Switzerland indicate the presence of trade networks along the Danube River. The low number of connections to nearby regions in today’s Czech Republic, in contrast, might suggest a preference for specific trade routes, possibly along major rivers like the Danube, which facilitated broader regional exchange. The relevance of the Danubian network during the EBA has long been recognized archaeologically and was termed the “Danubian Sheet Circle” (“Danubischer Blechkreis”) (**Ruckdeschl 1978**) and the correlation of relatedness in genetics and material culture is relevant to note. However, this relationship existed only during the EBA and there is a marked discontinuity in the Lech Valley between the EBA and the Middle Bronze Age (MBA) with regard to both material culture as well as social systems, which might point to the arrival of new groups in the Lech Valley from ca. 1700 BCE onwards (**Mittnik et al. 2019**). The genetic link between EBA Lower Austria and MBA Lech Valley is, therefore, unexpected. Thus, our findings might indicate that the population in the MBA Lech Valley likely originated from a group with strong connections to EBA Austria. It is important to note that Identical-by-Descent (IBD) links do not necessarily represent direct migration, but may indicate indirect links through a third, unrepresented region.

The examination of burial practices within this highly localized context also revealed intriguing insights. Previous studies have consistently identified patrilocality as a prevailing societal pattern in EBA Central Europe. However, this research uncovers instances of females of reproductive age who were buried with their fathers or brothers, along with their own offspring, suggesting that they did not marry into another community or were returned to their ancestral burial ground after death. This observation suggests the potential for studies of more nuanced female mobility and varied familial arrangements, ideally involving the study of strontium isotopes, which will enrich the existing interpretative framework and challenge dominant paradigms of patrilocality.

The examination of infants from Unterhautzental reveals the challenges faced by Early Bronze Age children. Three related toddlers from the same family all died at a young age, highlighting the harsh conditions Bronze Age children encountered early in their lives. Traces of injuries and diseases such as meningitis and pleural infection in the skeletal remains of these young individuals (**Rebay-Salisbury et al. 2018**) further illustrate the numerous challenges to health and survival within these communities. This fits well with the previously published case of child murder in Schleinbach (**Rebay-Salisbury et al. 2020**).

The extensive dataset, made possible by the dense sampling of individuals from burial grounds, provides a unique perspective on the Early Bronze Age communities in this region. It reveals the complexities of genetic relationships, significant exchange between groups, kinship and social structures. The findings shed light on population dynamics and emphasize the importance of microscale investigations to complement broader regional studies. This invites further exploration into the nuances of human interactions and sociocultural developments during this transformative period in history.

## Material and Methods

We collected 188 skeletal elements initially assigned to 188 ancient individuals from five EBA sites in Lower Austria described in SI Notes 1. Some samples have previously been analysed for mitochondrial DNA to access mother-chid relations (**Rebay-Salisbury et al. 2023**). All the samples were either teeth or petrous bones. The sampling procedures were conducted within a dedicated aDNA laboratory at MPI-EVA in Jena, following the laboratory’s archived protocols (https://doi.org/10.17504/protocols.io.bqebmtan and https://doi.org/10.17504/protocols.io.bdyvi7w6). In the case of complete crania from the collection of the Natural History Museum Vienna bone powder from the petrous part of the temporal bone was collected with a mobile sampling kit at the museum.

Samples were processed using an automated library protocol, which constructs libraries from single-stranded molecules. Each extract produced at least one library that was sequenced at a low depth on an Illumina HiSeq400 platform. The raw FastQC files underwent processing through the EAGER pipeline for adaptor removal, mapping against the human reference hs37d5, and PCR duplicate removal. The resulting information about library complexity and endogenous DNA percentage was combined with mapDamage estimates to assess the preservation of endogenous aDNA. To generate data from a large number of individuals, aDNA enrichment methods, using in-solution hybridization enrichment data consisting of approximately 1.2 million ancestry-informative positions (1240K capture) were applied to samples with 0.1% human endogenous DNA or more. Following the 1240K enrichment, the selected libraries were sequenced at standard depth (∼20 million reads). Post-1240K capture data was evaluated using EAGER and mapDamage with the same settings. All sequencing data from the same library or multiple libraries from one DNA extract or individual produced with the same protocols were processed equally and merged at the level of bam files. The authentication of aDNA involved three different methods, evaluation of aDNA damage with *mapDamage* (**Jónsson et al. 2013**), and contamination estimated on mtDNA with *schumtzi* (**Renaud et al. 2015**) and X chromsomes with ANGSD (**Rasmussen et al. 2011**), on the bam files to estimate modern DNA contamination on ancient samples. Genotypes were extracted from the pileups of ss-library bams, taking care to filter out bias due to damage.

Sampling of skeletal elements of 15 individuals (SI table 2) from the site Pottenbrunn was conducted in the dedicated ancient DNA clean rooms of of the Pinhasi Lab and later DNA was extracted DNA in the ancient DNA clean lab at Havard Medical School using a method specifically designed to retain short molecules (**Dabney et al. 2013, Rohland et al. 2018**) and created double-stranded DNA libraries (**Meyer and Kircher 2010, Kircher et al. 2012**). To minimize characteristic base misincorporations that accumulate in ancient DNA strands (**Rohland et al. 2015**), all libraries were partially treated with Uracil-DNA Glycosylase (UDG). For four libraries, we employed an in-solution hybridization approach to enrich the libraries for approximately 1.2 million single nucleotide polymorphisms (‘1240K’ SNP set) (**Fu et al. 2013, Haak et al. 2015, Mathieson et al. 2015**). For the remaining eleven libraries, we utilized the ‘Twist’ Ancient DNA capture protocol, which targets around 1.35 million SNPs—largely overlapping with the 1240K SNP set—using only a single round of enrichment (**Rohland et al. 2018**). Sequencing was performed on an Illumina NextSeq500 instrument using v2 150 cycle kits for 2 × 76 cycles and 2 × 7, or on a HiSeq X10 using v2.5 kits for 2 × 100 cycles and 2 × 7 cycles.

Following data merging and quality control, our final dataset was merged with publicly available genotype datasets of ancient and modern individuals from across Eurasia. We performed population structure analysis using PCA with modern West Eurasian populations as a reference and projected ancient individuals onto this PCA to avoid bias introduced by high rates of missing data in aDNA.

Relative proportions of ancestry components in the newly sequenced individuals were estimated using *qpAdm* (version: 632) from ADMIXTOOLS (**Patterson et al 2012**, https://github.com/DReichLab) using a threshold of 100k SNPs for analysis on an individual level (Supplementary Note 3) and modern reference individuals (Mbuti, Papuan, Han, and Karitiana) from the HO dataset and published ancient individuals (Ust Ishim, Villabruna, MA1).

We examined SNPs encoding for biological traits, such as LP, and eye/skin pigmentation, following the list of SNPs used in SI table 6. For each phenotype-associated locus, we report the number of reads with derived alleles versus the total number of reads covered on this site in SI table 6, by applying *SAMtools* pileup on BAM files after quality filtering (-q 30 -Q 30).

For the identification of closely related individuals, we employed the READ method. This approach estimates the coefficient of relatedness between two individuals by calculating the rate of mismatching alleles (P0) normalized with the pairwise allele differences among unrelated individuals within the population (α). This normalization corrects for SNP ascertainment, marker density, genetic drift, and inbreeding.

To detect relatives at a more distant degree, we utilized the *lcMLkin* with the options -l phred. This method leverages a maximum likelihood framework to infer identical by descent (IBD) on low-coverage DNA sequencing data from genotype likelihoods computed with bcftools. The coefficient of relatedness (r) is then calculated as k1/2 + k2, where k1 and k2 represent the probabilities of sharing one or both alleles IBD, respectively. The method can differentiate between parent–offspring (k0 = 0) and siblings (k0 ≥ 0, depending on the recombination rate. To ensure no bias between data generated with the 1240k and the Twist capture we compared pairwise mismatch rates within data of one capture and between the two captures (SI Note 2). Reconstructed family trees for all sites can be found in SI Note 4. This analysis was repeated using only SNPs on the X chromosome, as well as for the entire genome, utilizing the software *ngsRelate* (Korneliussen et al 2015).

To estimate Runs of Homozygosity (ROH), we employed the hapROH tool, which is specifically designed for inferring ROH in ancient DNA samples with pseudo-haploid data (**Ringbauer et al 2021,** SI Note 3). Genotype likelihoods were called using the MLE function of ATLAS, an ancient-DNA-specific caller (https://bitbucket.org/wegmannlab/atlas/). This was done across approximately 20 million SNPs from the 1,000 Genomes Project SNP panel, and these likelihoods were used as input for imputation through GLIMPSE. For imputation, we referenced phased haplotypes from the 1,000 Genomes Project phase-3 data and ran GLIMPSE with default settings, using sex-averaged genetic maps from HapMap as recommended (**Rubinacci et al 2020**). Haplotype IBD analysis was then carried out using ancIBD, a recently developed method that addresses the high error rates in phasing ancient DNA (**Ringbaur et al 2024**).

## Contributions

K.R.-S., and P.W.S conceived of the study. A.F., R.B., T.S., L.S., R.R., R.B., N.R., K.S., N.W., E.C., F.Z., K.C., L.I., L.Q., O.C., A.W., and G.B.M. performed laboratory work. A.F., G.U.N., A.M. and H.R., and analyzed data. K.R.-S., F.K., M.S., M.T.-N., F.N., and D.V. performed anthropological assessments. K.R.-S., F.K., M.S., M.T.-N., F.N., R.P. and D.V. assembled and interpreted archaeological material. A.F. and A.M. designed figures. A.F., A.M., and K.R.-S. wrote the paper with input from all co-authors.

## Acknowledgements

We thank the staff of the Department of Anthropology at the Natural History Museum in Vienna, in particular Karin Wiltschke-Schrotta, Doris Pany-Kucera and Michaela Spannagl-Steiner, for granting access to the human remains under their curation. The individuals can be found using the accession code NHMW-Anthro-OSTE and the inventory number (Inv. Nr.). We are grateful to Kurt Fiebig, the director of the excavation at Drasenhofen, for the detailed excavation records. Walther Parson and David Reich generously let us access and re-analyze samples and data from previous projects. This study was funded in the framework of the project “The value of mothers to society: responses to motherhood and child rearing practices in prehistoric Europe’, which received funding from the European Research Council (ERC) under the European Union’s Horizon 2020 research and innovation programme (grant agreement No 676828, PI: K. Rebay-Salisbury).

