## Supplemental Information for "Tracing social mechanisms and interregional connections in Early Bronze Age Societies in Lower Austria"

**Supplementary Information: “Genetic Perspectives on Early Bronze Age Societies in Lower Austria: Tracing societal mechanisms and Interregional Connections”**

### SI Note 1: Archaeological Information on the sites

#### Drasenhofen

Archaeological excavations on both sides of the Stützenhofner Bach creek were initiated in 2018 in response to the construction of a motorway bypass for Drasenhofen at the Austrian-Czech border (Fiebig and Csaplaros 2019). The Early Bronze Age site includes extensive settlement remains, dating to the classical Únětice culture and early Vĕteřov culture (c. 2150-1700 BC). Four isolated buildings, consisting of two workshops and two dwellings, were found at the southeastern end of the excavation area, forming a small farmstead separated from the larger, village-like settlement (Fiebig and Csaplaros 2019). The cemetery was located south of the village and north of the farmstead, with 25 burials in 22 graves arranged in four rows potentially representing former inhabitants of the settlement (Horváth 2019, Kanz 2019). The topography of the site suggests that all the graves of the cemetery were located within the construction line and the grave group was completely excavated. The cemetery's block-like, regularly rectangular shape of approximately 30 × 50 m implies that its boundaries were likely demarcated, for example by a fence. Outside the cemetery, four additional individuals were discovered buried in former storage pits within the settlement area. With 15 female and 14 male individuals, the sex ratio is balanced; eleven individuals died before they reached the age of 20. In the course of our investigation, we found the oldest genetic evidence of two different strains of the plague bacteria (Yersinia pestis) in male individuals, who died at the age 23-30 and 22-27 years of age, respectively. To date, this is the earliest evidence of Late Neolithic / Early Bronze Age plague in today’s Austria (Neumann *et al.* 2023).

#### Zwingendorf

In 1977, the archaeological site of Zwingendorf, Austria, yielded a total of nine Early Bronze Age graves during excavation in 1977 (Grefen-Peters 1982, Wewerka 1982). A radiocarbon sample from Zwingendorf (NHMW-ANTHRO-OSTE-1008585/8A) was sent to the Higham laboratory at the University of Vienna and measured at the Vienna Environmental Research Accelerator AMS facility. The results ranging from 2120 to 1945 cal. BC confirmed the typo-chronological dating that has placed the site within the developed Únětice culture, estimated from approximately 2000 to 1600 BC. The small grave group with north-south orientated grave pits revealed ten remarkably undisturbed individuals interred in flexed positions, all on the right sides of their bodies with their heads towards the south. Grave 8 included a 4-year-old child with a bronze ring placed in front of a 15-16-year-old adolescent’s body. The grave further contained five ceramic vessels.

#### Unterhautzenthal

Unterhautzenthal, an Early Bronze Age site of the Únětice culture, comprises both remnants of a settlement, including storage pits in which human remains have been found, and a small cemetery excavated in the 1980-90s (Lauermann 1995, Lauermann 1997, Lauermann, Pucher, and Schmitzberger 2001). Recent re-analysis of the human remains from Unterhautzenthal has provided evidence of additional individuals, primarily neonates and children, who were not individually documented at the time (Rebay-Salisbury *et al.* 2018). The inclusion of these skeletons has expanded the total count of excavated individuals at Unterhautzenthal to fifty-eight, with thirty-two (55%) representing sub-adults under the age of twenty. Almost all bodies were placed in flexed position on the right side of the body; one was found in prone position. The predominant body orientation was with the head to the south-west, south and south-east, with a few exceptional cases oriented to the north. The triple burial 95 contained the remains of a 35-45-year-old woman and two children, who died at the ages of 2-3 and 4-5 and were placed in close contact to the woman’s body. The burial of two closely related children in a storage pit, a 2 and a 6-year-old in embrace, is further noteworthy (Rebay-Salisbury *et al.* 2023).

#### Schleinbach

Schleinbach, situated 10 km northeast of Vienna in Lower Austria, lies north of the Danube and represents a cemetery and settlement complex associated with the Únětice culture. The Bronze Age settlement, encompassing approximately 1.5 hectares, likely neighbored a prehistoric lake. Initial discoveries were made in 1911 within a brick factory, and more or less systematic excavations accompanied the clay extraction until the 1980s (Rebay-Salisbury *et al.* 2020, Rettenbacher 2004). Within the site, two distinct groups of Early Bronze Age burials were uncovered: Group 1 located in the western area and Group 2 in the eastern area of the site, at approximately 180 m distance. The western grave group with graves arranged in two rows included the double burial of two male individuals 30/31 with identical head fractures (Weninger 1954). Storage pits with human remains, such as the quadruple burial of an adult male and three children in Pit 60 (Weninger 1954) and the burial of a 5-6-year-old boy with no fewer than four fatal blunt force traumata to the skull in Pit 3/1981 (Rebay-Salisbury *et al.* 2020) contribute to the richness of the site's historical record and its recognition within Austrian prehistory. The eastern grave group included 17 burials, many of which were heavily disturbed, with few grave goods and intermingled human remains. The skeletal remains of 62 individuals were available for a recent osteological re-analysis (Pany-Kucera *et al.* 2020), which found a demographic composition of 14 adult and mature females, 15 adult and mature males and 27 subadults, some of which were fetuses and neonates represented only by a few bones. The study further found extraordinary levels of stress indicators and traumas at the site, testifying to violence, conflict, abuse and marginalization at Schleinbach.

#### Ulrichskirchen

Ulrichskirchen is an Únětice culture site complex excavated in 2011 in advance of a pipeline construction, with settlement traces, including depositions of human remains, and a group of graves. The Schleinbach and Ulrichskirchen sites are only about 1km apart, which allows them to be understood as one larger, contemporary complex. The first archaeological intervention revealed a small cemetery comprising 12 individuals and five further individuals from former storage pits, which are currently prepared for publication by Maria Teschler-Nicola and Friederike Novotny. The second archaeological intervention led to the discovery of six further individuals buried in storage pits, which were subject to Domnika Verdianu’s Master thesis (Verdianu 2024). The graves were arranged in two rows of five graves each, and a further row of two graves; the archaeological intervention, however, only covered the width of the pipeline, so further graves can be expected. All graves were aligned along the north-east / south-west axis and contained single burials. The bodies were placed head south on the right side in flexed position, and all graves showed traces of grave re-opening some time after the burial. One settlement pit contained four individuals, one two individuals, and the others a single individual each. Whereas the multiple depositions in particular make the impression that bodies were thrown in and discarded, three single depositions seem carefully placed, one of which even oriented according to the prevailing funerary customs.

#### Franzhausen

The cemeteries of Franzhausen I and II, situated in the Traisen Valley of Lower Austria, are part of an expansive archaeological landscape revealed by large-scale rescue excavations in the last decades of the 20^th^ century. The ca. 2200 burials span the entire period of the Early Bronze Age from about 2300–1600 BC and are associated with the Unterwölbling cultural group (Neugebauer 1994). The dead were buried in flexed, gender-specific body position, usually in individual graves. Women were placed on the right side of the body, head south, whereas men were placed on the left, head north. Grave goods comprise bronze costume and jewellery, weapons and tools, ceramic vessels as well as meat produce. Merely twelve of the 716 excavated graves at Franzhausen I contained double or multiple burials (Neugebauer and Neugebauer 1997: 26). The triple burial 599 is one of these exceptional cases, where the remains of a 20–25-year-old male individual (designated as 599A) were found positioned on the left side with the head oriented to the north-west in a wooden coffin. The upper part of the man's body was severely disturbed and his cranium was missing. Two adolescent individuals, 14–16 (designated as 599B) and 12–14 (designated as 599C) years at the time of death, accompanied the adult male. Their bodies were positioned at the adult’s feet in the south-eastern part of the coffin, on their left sides with their heads to the south-east. The lower legs of the young individuals came into close contact, suggestive of a deliberate placement. The burial sequence indicates that the adult male was laid in the coffin first, followed by the older adolescent, and finally, the youngest was positioned behind the older one's back in a single act of deposition (Rebay-Salisbury 2018).

#### Pottenbrunn

The site of Pottenbrunn (Blesl 2006, Novotny 2006) is situated in Lower Austria south of the Danube in the Traisen Valley, approximately 12 km south of the more famous sites Franzhausen I and II. The site was excavated in 1981/82 in the course of rescue excavations, following the same digging and recording system as for Franzhausen. Pottenbrunn belonging to the Unterwölbling culture seemed to have been completely excavated and includes Early Bronze Age settlement structures, with graves placed in abandoned areas of the settlement, and a total of 74 roughly north-south oriented grave pits. Of those, 69 included human remains. All were single graves, but four graves included remains of more than one individual (Blesl 2006: 18). The site was heavily disturbed, not only by contemporaneous grave re-opening, but also by a La Tène cemetery built on top of the site several hundred years later (Ramsl 2002). The cemetery population includes 19 females and 20 males, as well as 37 subadult individuals (Novotny 2006). In many ways, burial practices are comparable to Franzhausen I, with gendered burial placement, orientation and grave goods, but overall poor preservation.

### SI Note 2: Evaluation of bias from different wet-lab methods on genetic kinship analysis

Of the 15 samples processed in Boston, 11 (I28636, I28642, I28991, I28992, I29242, I29247, I29687, I29691, I29693, I11700, I11704, I11705, I28635, I28641, I29246) were captured using a new enrichment method (Twist1.4M).

To ensure that combining these samples with those captured using the 1240k method would not introduce bias, we analyzed pairwise mismatch rates (Fig. 1). While there is a slight tendency for the 1240k samples to be more similar to each other than to the Twist samples, this bias is minimal and does not affect the relatedness outcomes in our relationship analysis.
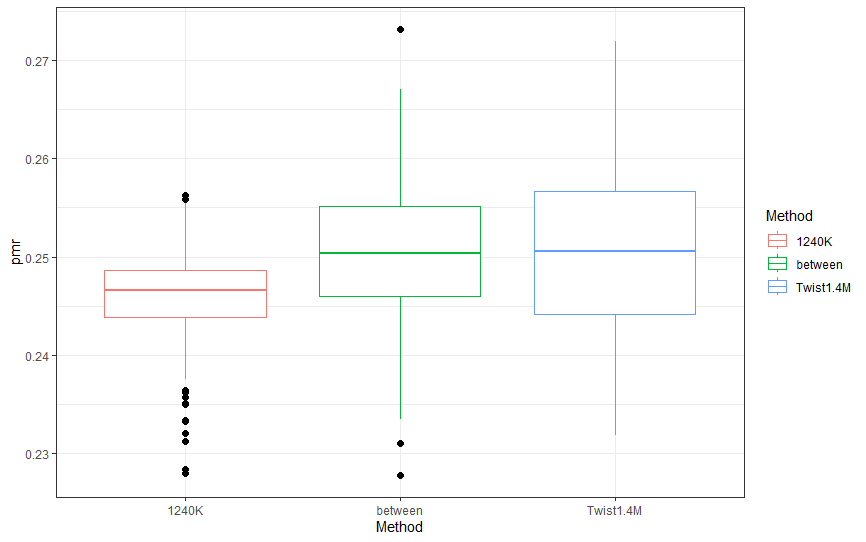

**Figure 1: Pairwise mismatch rates within 1240k and Twist samples (red and blue) and between the two groups (green).**

### SI Note 3: Analysis of runs of homozygosity

We used the hapROH method (v.1.0) for the analysis of runs of homozygosity (ROH), which is specifically designed to analyze low-coverage aDNA data. By leveraging linkage disequilibrium from a panel of modern haplotype references, this method can successfully infer ROH longer than 4 cM on 1240K data with at least 0.3× coverage. In cases of close parental relatedness, which results in long ROH in offspring, hapROH efficiently detects very long ROHs even at lower coverage. We called ROH individuals with >250,000 SNPs. The program's embedded functions were used for plotting the ROH as combined histograms.

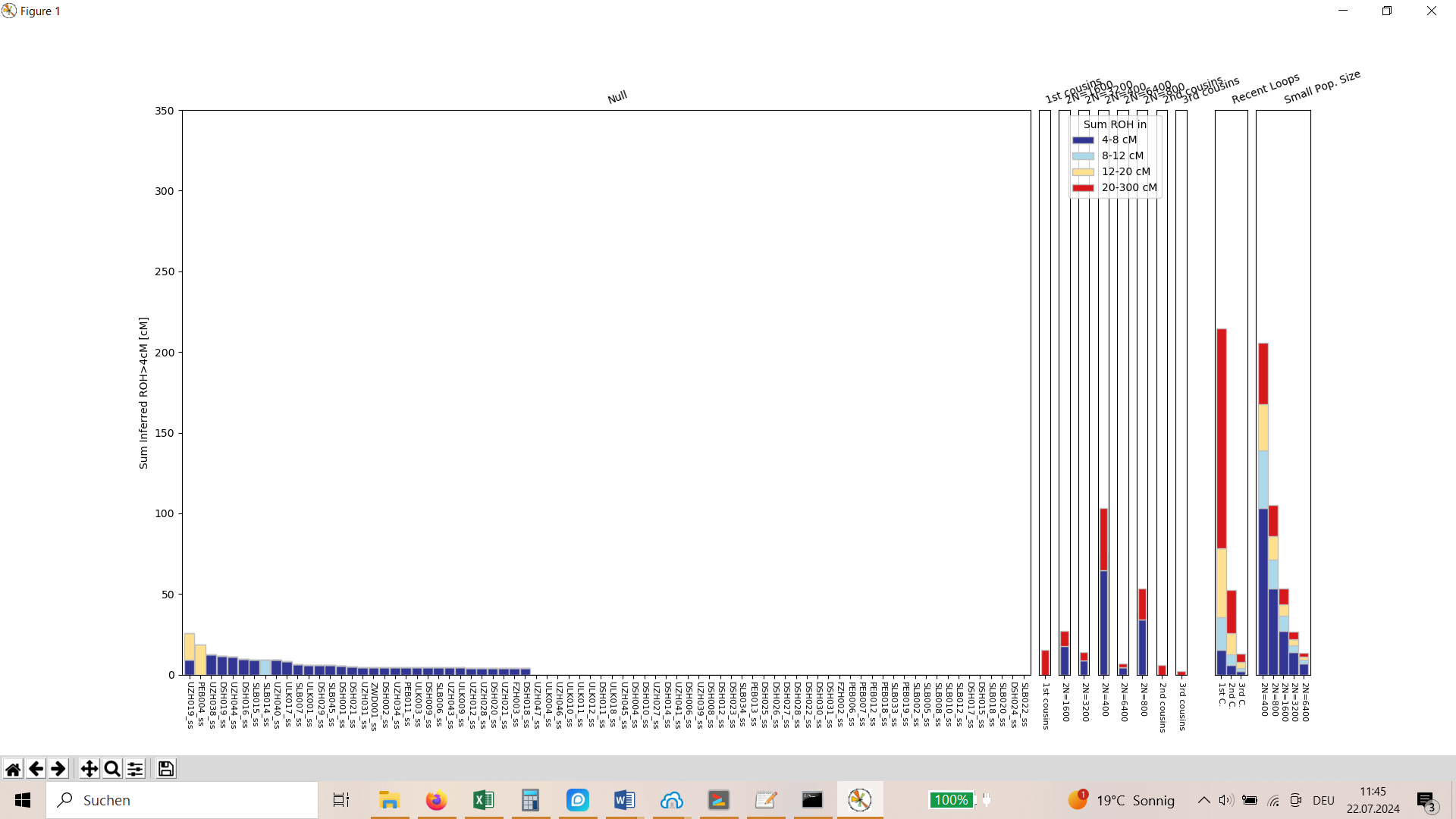

**Figure 2: Sum of runs of homozygosity (ROH) per individual.** Left the individuals from the EBA sites in Lower Austria and right expected patterns of ROH for cousin-cousin marriage and small population size.

### SI Note 4: Reconstructed family trees per site

#### Drasenhofen

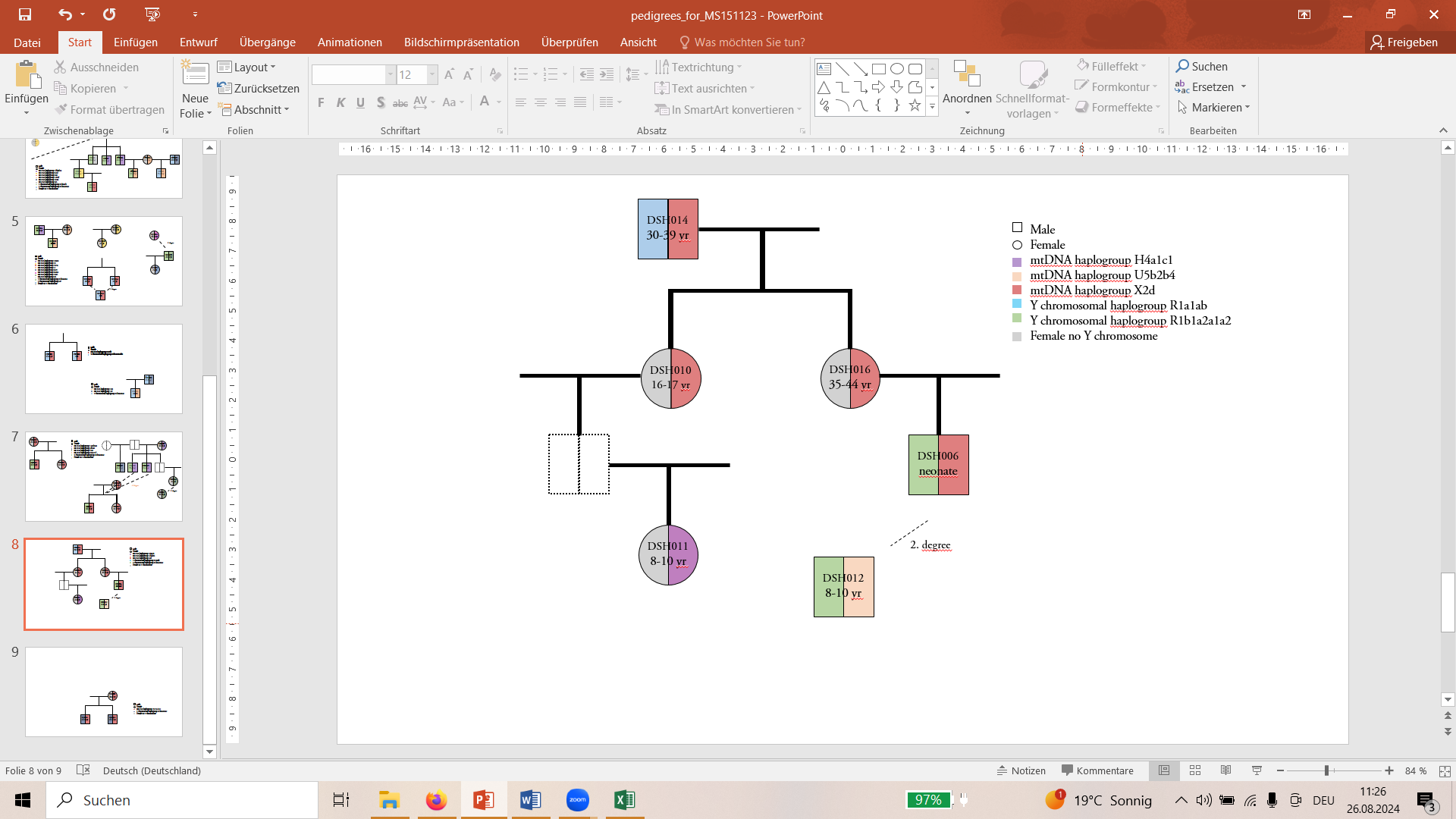

**Figure 3: Reconstructed family tree 1 Drasenhofen.**

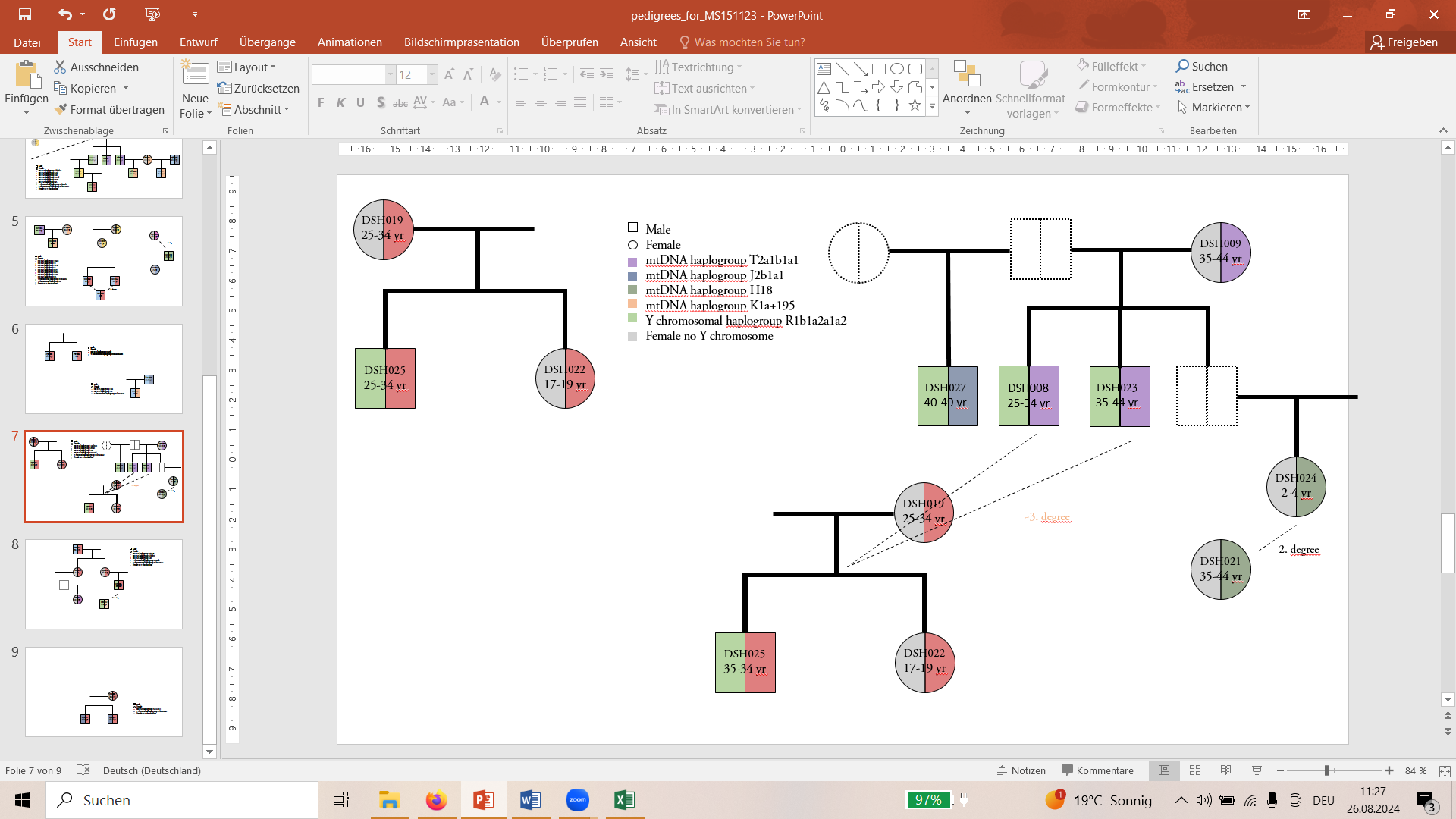

**Figure 4: Reconstructed family tree 2 Drasenhofen.**

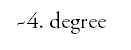
In addition to the family connections shown, the following individuals are more distantly related: DSH006 is approximately 4^th^ degree related to DSH008, DSH023 and DSH025. DSH012 is approximately 5^th^ degree related to DSH023 and DSH008. Approximately 6^th^ to 8^th^ degree related is DSH009 to DSH006, DSH014 and DSH016.

#### Zwingendorf

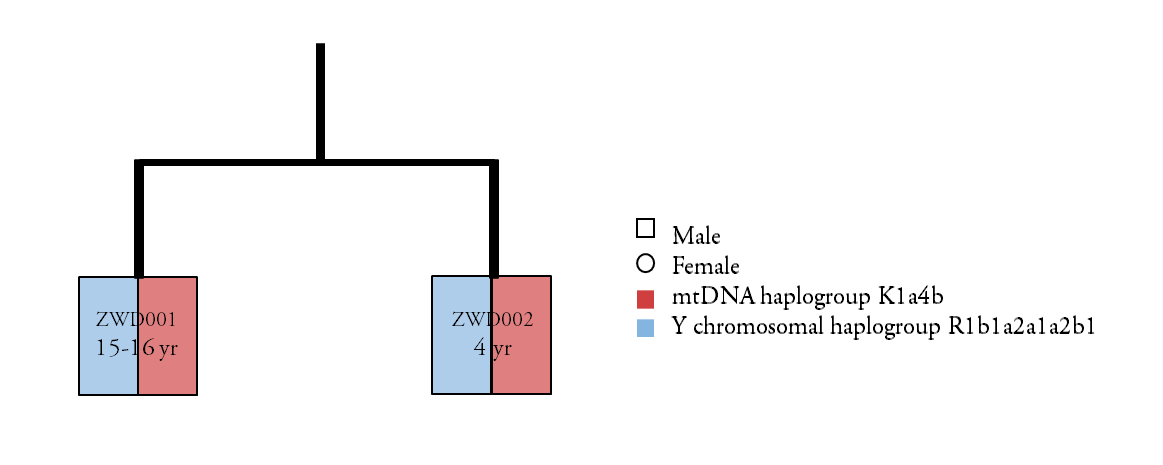

**Figure 5: Reconstructed family tree Zwingendorf.**

#### Unterhautzental

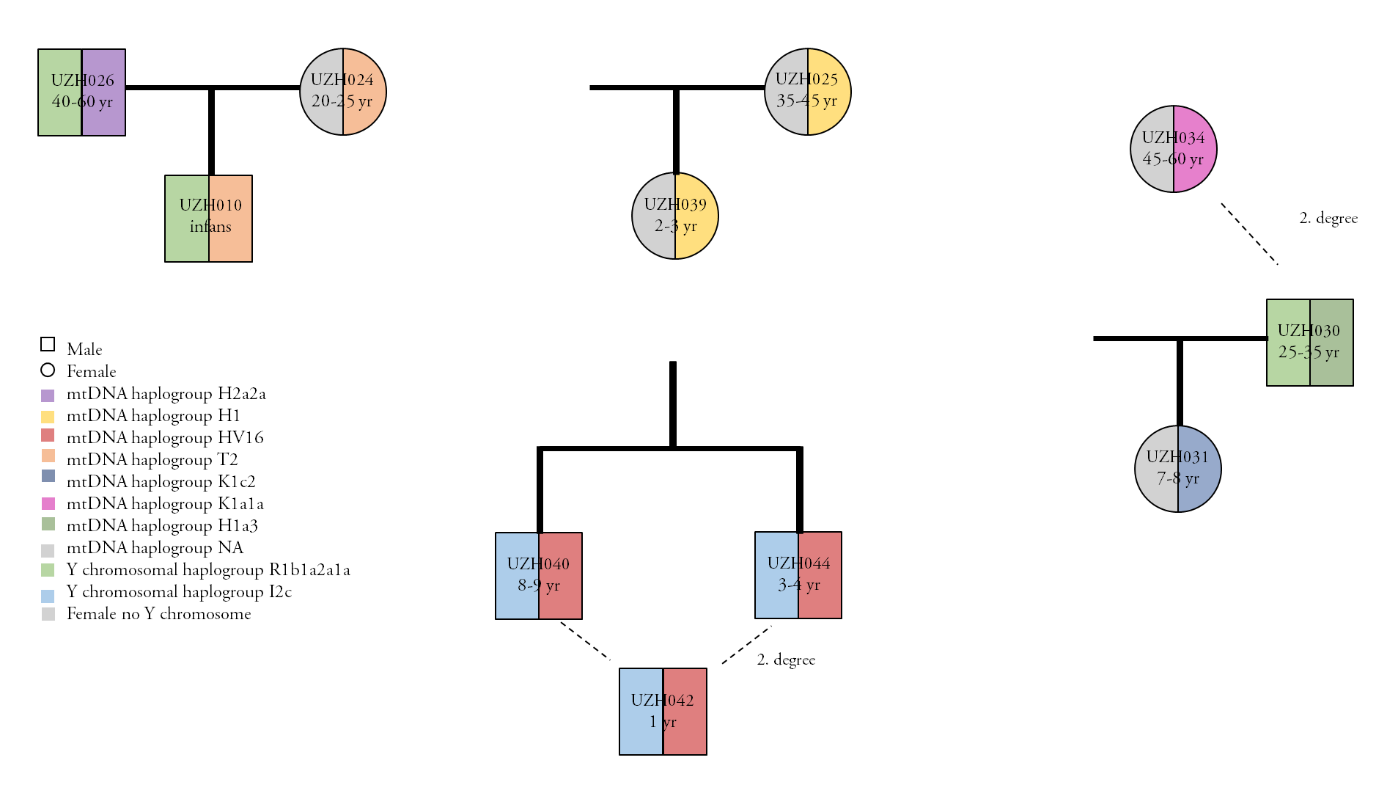

**Figure 6: Reconstructed family trees Unterhautzental.**

#### Schleinbach

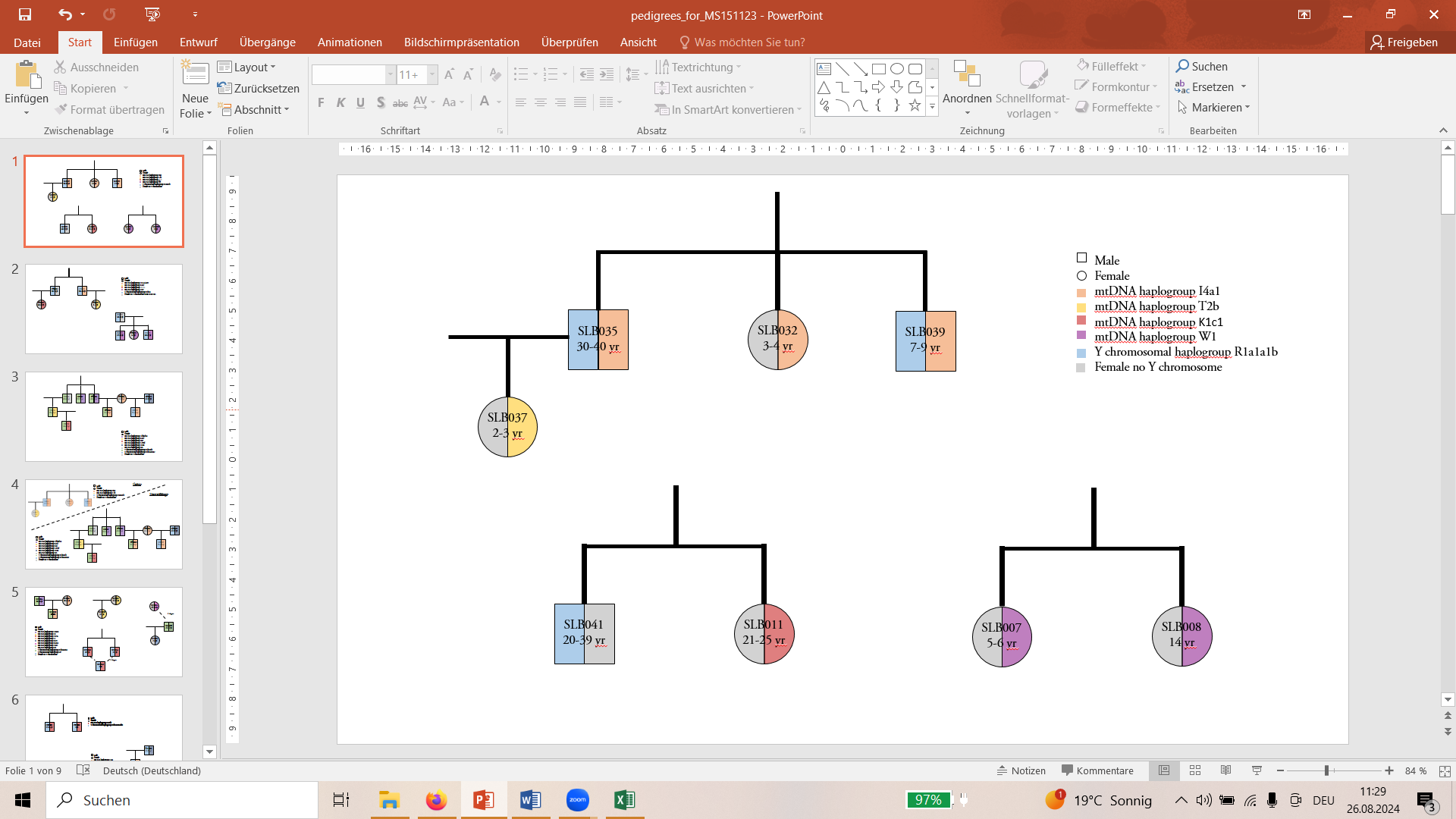

**Figure 7: Reconstructed family trees Schleinbach.**

#### Franzhausen

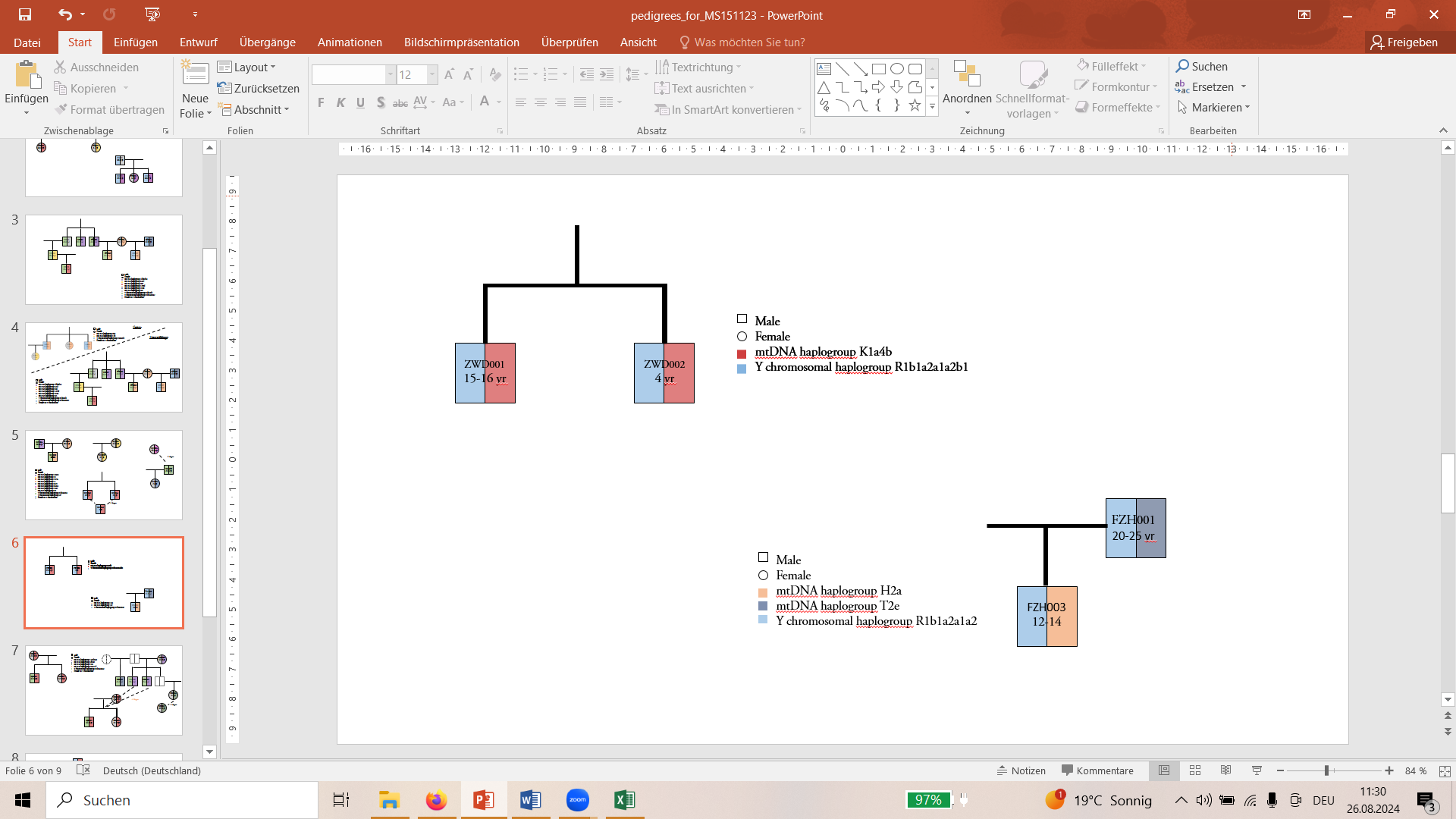

**Figure 8: Reconstructed family tree Franzhausen.**

#### Pottenbrunn

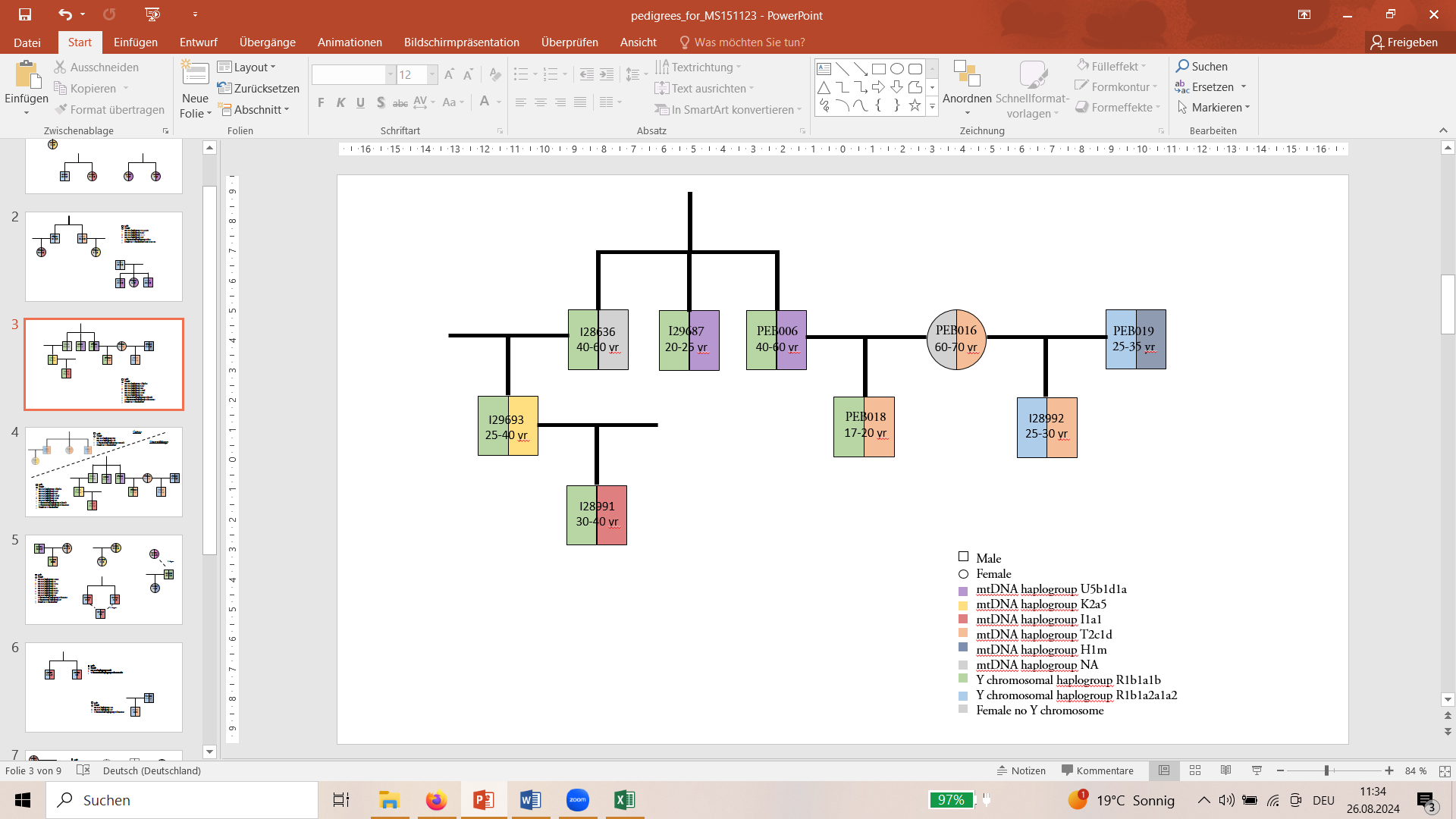

**Figure 9: Reconstructed family tree 1 Pottenbrunn.**

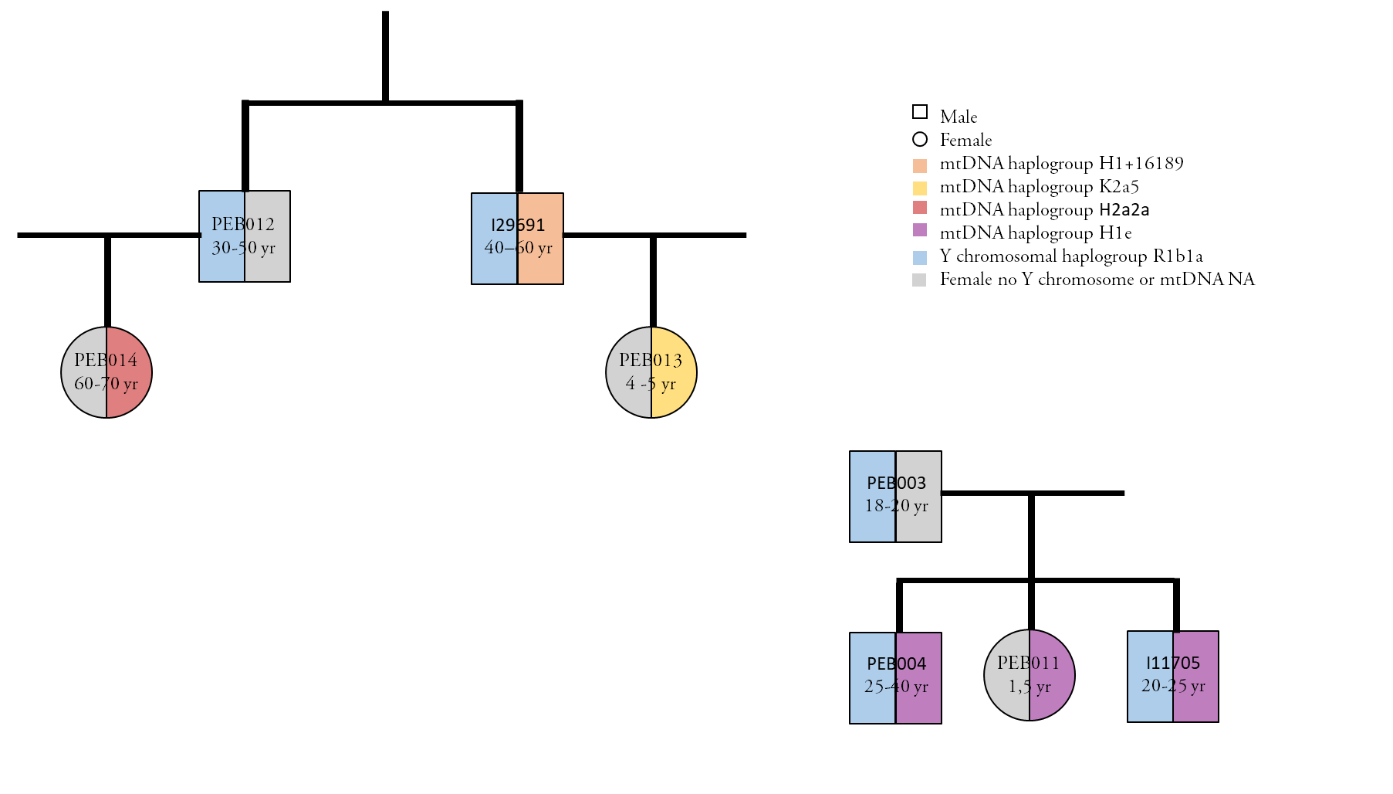

**Figure 10: Reconstructed family tree 2 Pottenbrunn.**

#### Ulrichskirchen

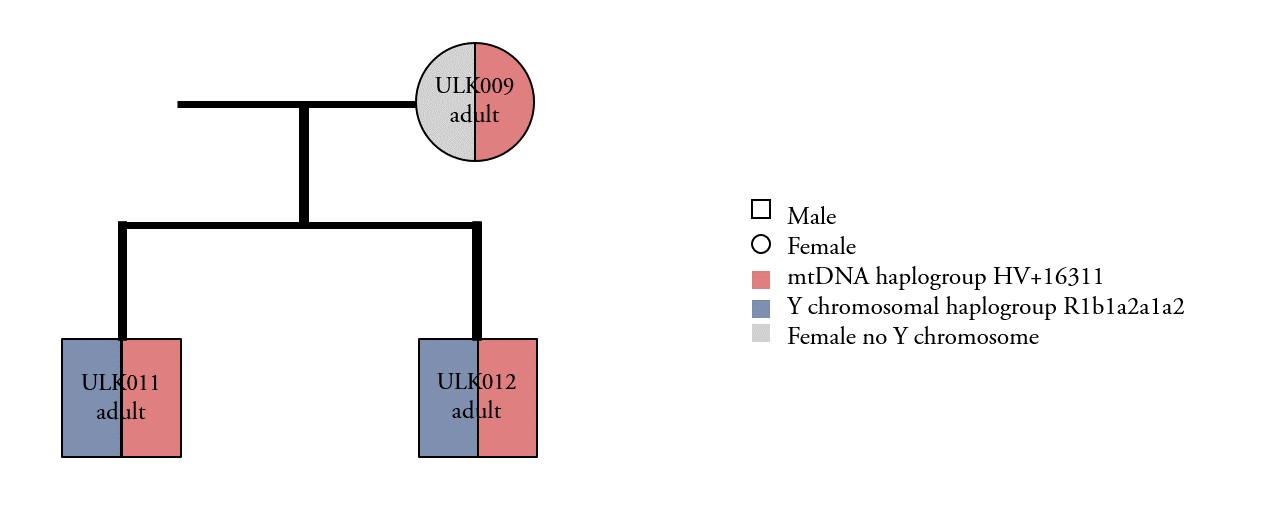

**Figure 11: Reconstructed family tree Ulrichskirchen.**

### SI Note 5: Separate IBD networks for males and females

IBD networks were constructed separately for males and females to assess differences in their levels of connectedness. To avoid bias, children for whom parental relationships could be identified in the dataset were included in the analysis. This was done to ensure that the number of connections observed for any individual was not artificially inflated or reduced based on known family ties, which could skew the comparison between males and females. The results show that males generally have more connections than females. Additionally, there are more females without any connections compared to males—five versus two in the Únětice culture and one versus none in the Unterwölbling culture. This pattern further supports the idea of a patrilineal society in these groups.

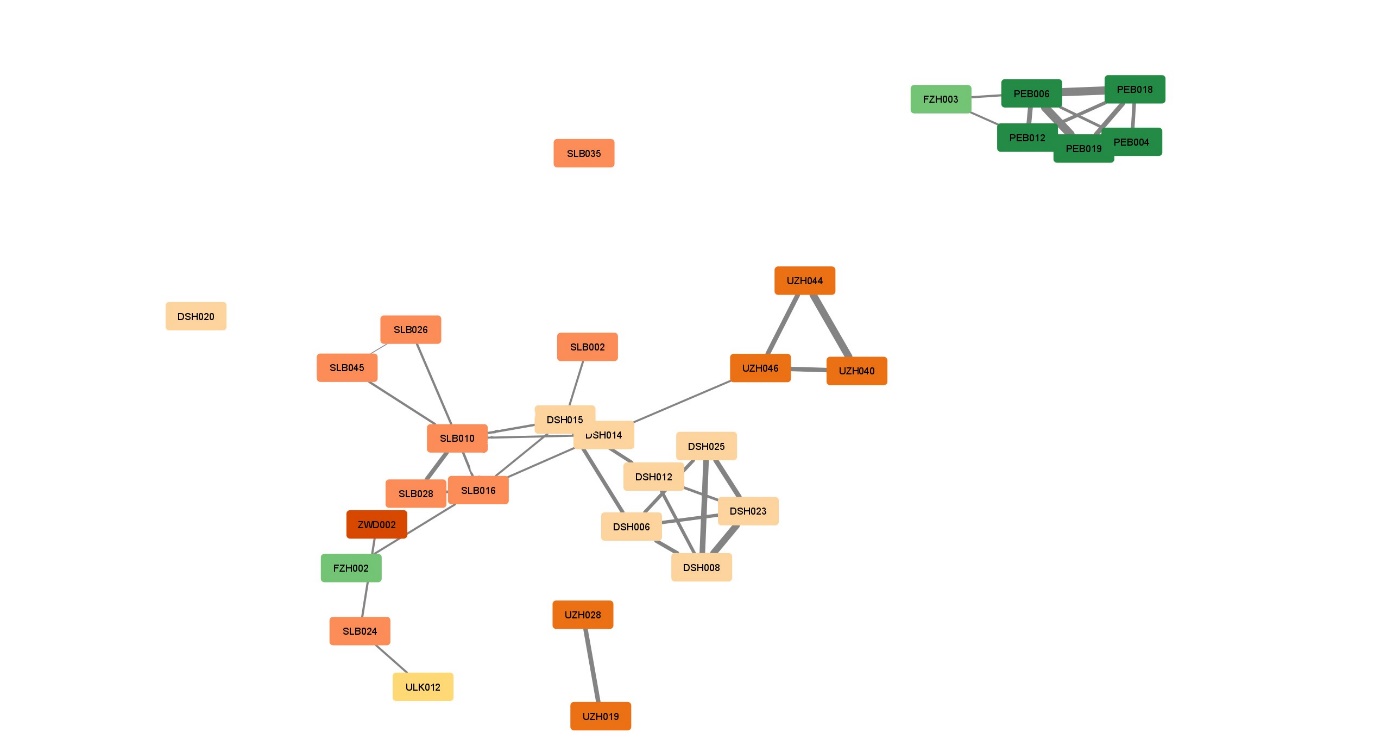

**Figure 12: IBD network for males in the Únětice (orange) and Unterwölbling (green) cultures.** Each square represents an individual male, with different shades indicating the specific site within each culture. The strength of the lines between individuals reflects the maximum IBD (Identity by Descent) value, with thicker lines representing stronger genetic connections.

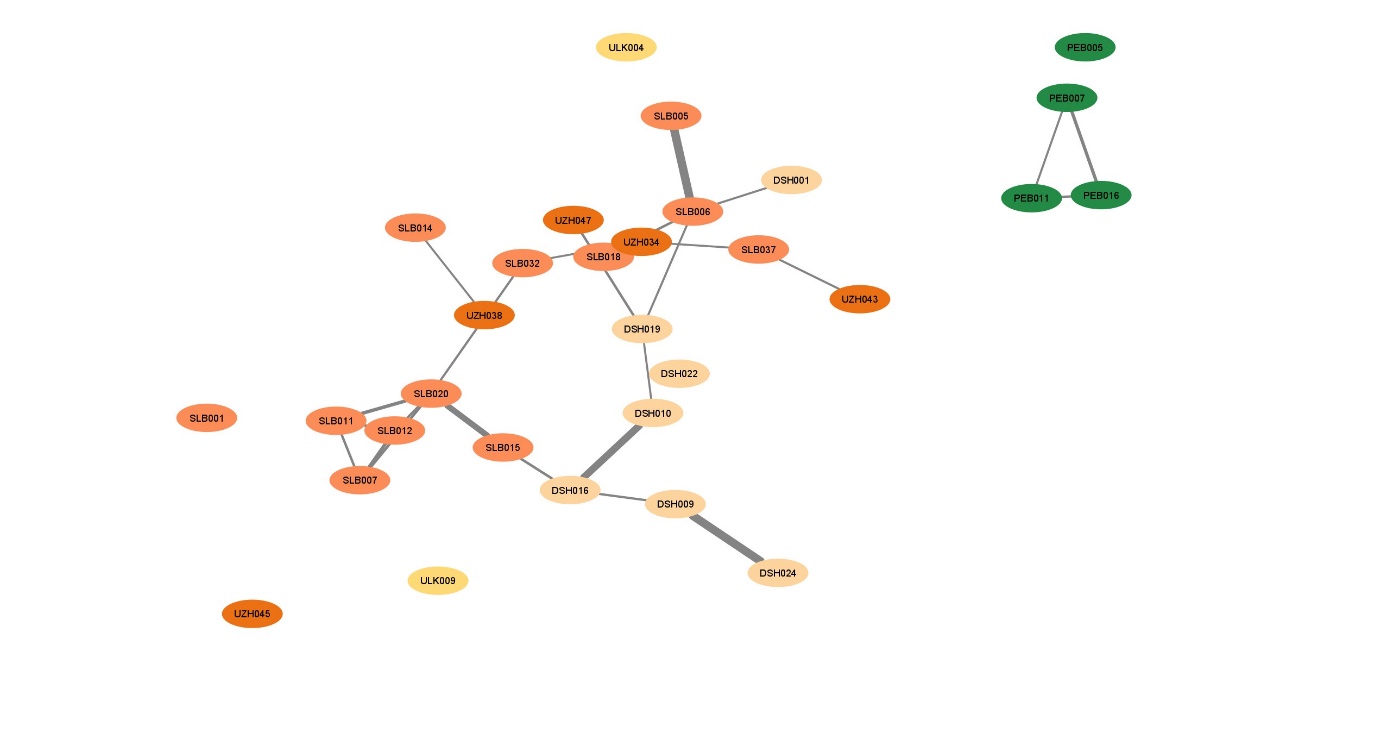

**Figure 13: IBD network for females in the Únětice (orange) and Unterwölbling (green) cultures.** Each circle represents an individual female, with different shades indicating the site within each culture. As with the male network, the strength of the connecting lines indicates the maximum IBD value, with thicker lines showing stronger genetic relationships. The network shows fewer connections among females, with more individuals having no connections, especially in the Únětice culture. This pattern aligns with observations of gender-based differences in social structure and suggests a patrilineal society.
